# Constructing a multiple-layer interactome for SARS-CoV-2 in the context of lung disease: Linking the virus with human genes and co-infecting microbes

**DOI:** 10.1101/2021.12.05.471290

**Authors:** Shaoke Lou, Tianxiao Li, Mark Gerstein

**Affiliations:** Yale University

## Abstract

The severe acute respiratory syndrome coronavirus 2 (SARS-CoV-2) pandemic has caused millions of deaths worldwide. Many efforts have focused on unraveling the mechanism of the viral infection to develop effective strategies for treatment and prevention. Previous studies have provided some clarity on the protein-protein interaction linkages occurring during the life cycle of viral infection; however, we lack a complete understanding of the full interactome, comprising human miRNAs and protein-coding genes and co-infecting microbes. To comprehensively determine this, we developed a statistical modeling method using latent Dirichlet allocation (called MLCrosstalk, for multiple-layer crosstalk) to fuse many types of data to construct the full interactome of SARS-CoV-2. Specifically, MLCrosstalk is able to integrate samples with multiple layers of information (e.g., miRNA and microbes), enforce a consistent topic distribution on all data types, and infer individual-level linkages (i.e., differing between patients). We also implement a secondary refinement with network propagation to allow our microbe-gene linkages to address larger network structures (e.g., pathways). Using MLCrosstalk, we generated a list of genes and microbes linked to SARS-CoV-2. Interestingly, we found that two of the identified microbes, Rothia mucilaginosa and Prevotella melaninogenica, show distinct patterns representing synergistic and antagonistic relationships with the virus, respectively. We also identified several SARS-COV-2-associated pathways, including the VEGFA-VEGFR2 and immune response pathways, which may provide potential targets for drug design.

## Introduction

Severe acute respiratory syndrome coronavirus 2 (SARS-CoV-2) has caused one of deadliest pandemics in human history, infecting more than 175 million people and causing 3.7 million deaths (WHO, Jun 2021). Thus, there is an urgent need to understand the mechanisms governing viral infection and host responses in order to develop effective methods for diagnosis, treatment, and quarantine.

Based on in-depth investigations of the SARS-CoV-2 genome and transcriptome structure (Kim, Lee et al. 2020, Wu, Zhao et al. 2020), researchers have elucidated the SARS-CoV-2 infection pathway (Chen, Malone et al. 2020, Viswanathan, Arya et al. 2020), (Kim, Lee et al. 2020, Shu, Huang et al. 2020, Yin, Mao et al. 2020), (Schoeman and Fielding 2019) and have identified several infection-related pathways within the human host (Ho, Mok et al. 2021). These efforts have revealed key processes in SARS-CoV-2 infection and serve as the cornerstone for further large-scale regulatory network and biosignature studies.

High-throughput methods have also elucidated interactions between SARS-CoV-2 and the host, shedding light on the host protein/virus protein interaction network (Gordon, Hiatt et al. 2020, Gordon, Jang et al. 2020), perturbations in the host gene and cellular networks during the initial stages of SARS-CoV-2 infection (similar to the triggering of cytokine storms) (Li, Guo et al. 2021), and interactions between host proteins and SARS-CoV-2 RNA during active infection (Flynn, Belk et al. 2021). Single-cell RNA sequencing has also provided valuable information regarding biological pathways and biosignatures (Zhang, Wang et al. 2020, Ng, Granados et al. 2021) and has revealed the large-scale cellular and molecular landscape of immune responses during SARS-CoV-2 infection in multiple tissues (Guo, Li et al. 2020, Zhang, Wang et al. 2020, Ren, Wen et al. 2021).

Despite recent achievements, our current understanding remains insufficient to provide an integrative picture of the virus-host interaction. For example, research has shown that microRNAs (miRNAs) play an important role in antiviral immune responses (Mirzaei, Mahdavi et al. 2021) and participate in the host response to SARS-CoV-2 (Arisan, Dart et al. 2020, Jafarinejad-Farsangi, Jazi et al. 2020), and potential miRNA binding sites occur on the SARS-CoV-2 genome (Jafarinejad-Farsangi, Jazi et al. 2020). Moreover, Previous studies have shown that COVID-19 patients usually present with co-infection (Lansbury, Lim et al. 2020). The co-infecting microbes may have similar host responses and pathogenic effects and form co-abundant clusters. Numerous other microbes play an indispensable role in shaping the host immune response, but their effects on SARS-CoV-2 infection remain largely unknown. Also, a recent study reported differential bacterial compositions between COVID-19 patients and healthy controls, with COVID-19 patients having a lower diversity (Xu, Lu et al. 2021). This finding indicates that unknown interactions occur between upper respiratory and gut microbiomes during SARS-CoV-2 infection, suggesting their potential importance in the host response.

Overall, it is challenging to integrate these multiple layers of information, including gene, miRNA, and other microbes’ information, and to identify inter-layer associations relevant to the host response in SARS-CoV-2 infection. In this paper, we propose an advanced option, MLCrosstalk, for elucidating host-pathogen interactions. MLCrosstalk incorporates multiple data resources and features and identifies both common and COVID-19-specific host gene-microbiome interactomes in different tissues across human diseases. Using network propagation analysis, we further extend our study to the flow of host responses.

## Results

### The MLCrosstalk model and validation

Our MLCrosstalk has three major advantages for performing integration analysis of multiple-type data. This approach 1) takes advantage of the Dirichlet distribution of hyperparameter to handle sparse and noisy data, 2) enforces a unitary topic distribution for each patient/sample to facilitate linkage identification between different types of data, and 3) easily extends to multiple data types with missing samples allowed (see Figure 1 for workflow). In our study of COVID-19 datasets, MLCrosstalk extracted dimensionally reduced patterns to infer a comprehensive linkage between host protein-coding genes, noncoding genes (e.g., miRNA), and microbes. MLCrosstalk defined a comprehensive interactome for the gene-microbe-miRNA network. The network then underwent further refinement via network propagation to integrate pathway information and connect hostpathogen interactions with biological relevance.

**Figure 1.**
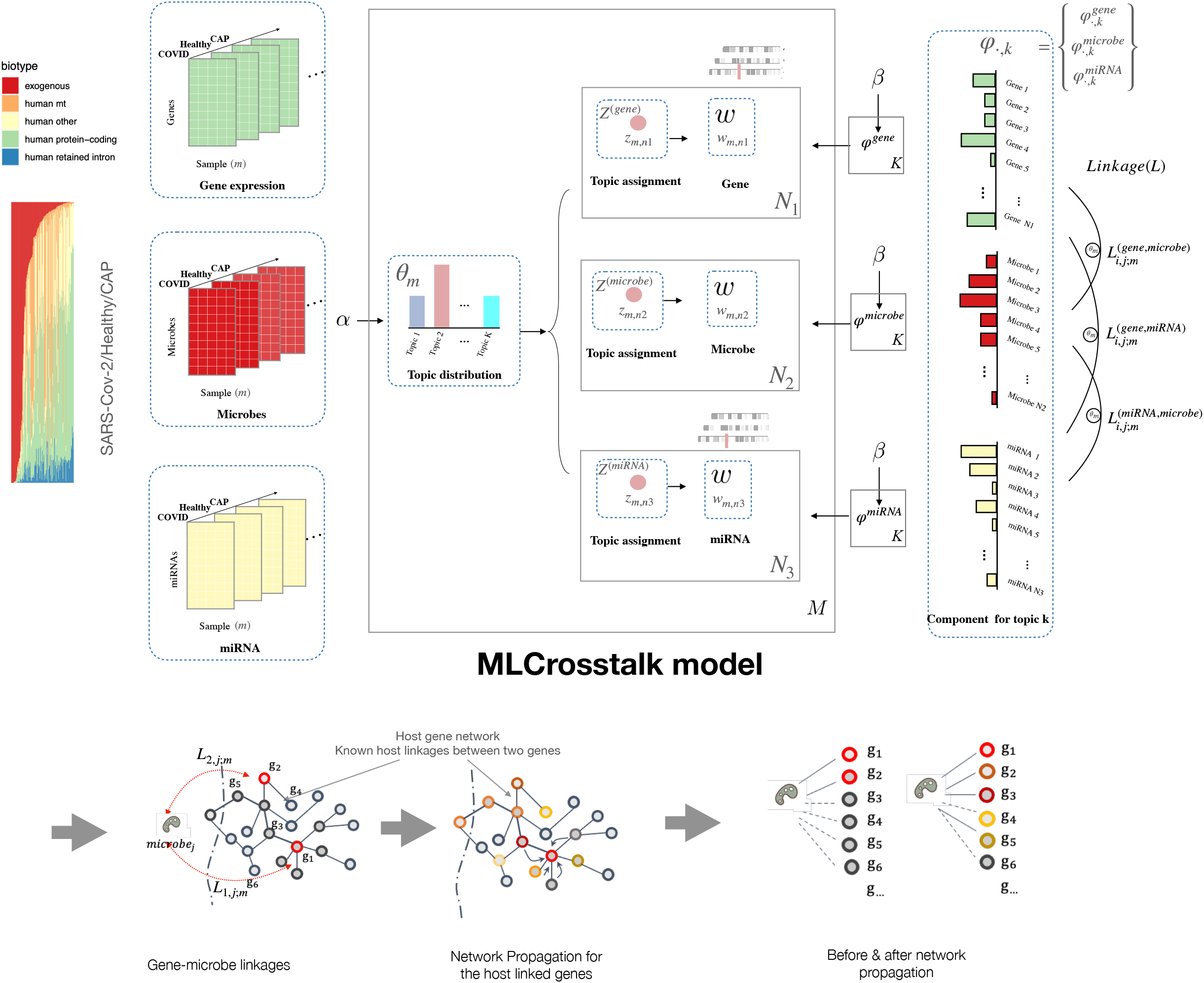
MLCrosstalk workflow. We transform gene expression, microbe abundance, and (pre)miRNA expression data, which are then input into the MLCrosstalk model. After training, we apply network propagation to refine the linkages. Multiple layer comparison and network tracing can identify shared and specific pathways and connections.

We first evaluated the trained model by exploring the clustering of topic distributions for the samples. The clustering results indicate that the topic distribution can capture the patterns across disease and tissue groups. The trained model groups most of the COVID-19, healthy, and community-acquired pneumonia (CAP) individuals into distinct clusters. Using the Kullback-Leibler divergence between topic distributions in comparison to a random background, thus, we found that Topic 9 differs the most from the random background distribution and is the most interesting topic.

We then annotated the functions of the top-weighted genes from Topic 9 using multiple known gene sets, including the Kyoto Encyclopedia of Genes and Genomes (KEGG), WikiPathways, virus-host protein-protein interaction (PPI), and COVID-19-related gene sets. We found that these genes are highly enriched in immune-related pathways and heat-shock response proteins (Fig. 2B). Figure 2C-D shows the top-weighted proteincoding genes, miRNAs, and microbes, with SARS-CoV-2 being one of the strongest contributors for Topic 9.

**Figure 2.**
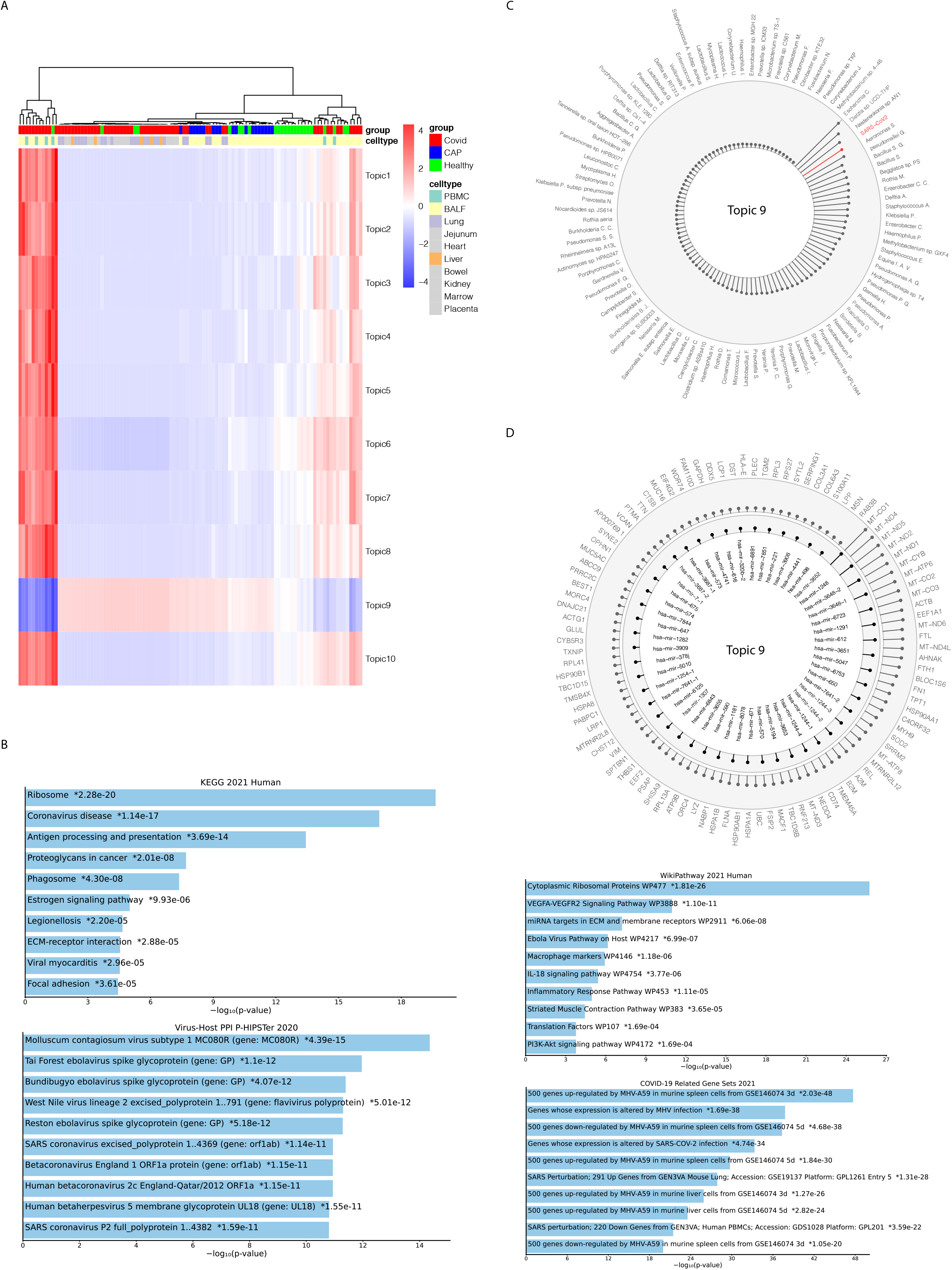
Model evaluation and functional analysis. A. Heatmap of the topic distribution across all SARS-CoV-2 samples. B. Functional analysis of the most interesting topic (Topic 9). C–D. The top-weighted protein-coding genes, pre-miRNAs, and microbes for Topic 9.

### SARS-Cov-2 links to Microbes

SARS-CoV-2 was the one of most common microbes among the COVID-19 patient samples. We investigated and compared the microbes that may be associated with SARS-CoV-2 based on the similarity from raw abundance, advanced feature weights (topic weights), and target functional linkage correlation. From the raw abundance of microbes, we took the top 100 most abundant microbes, and identified microbe groups based on the abundance.

We then used a different approach to identify COVID-19-associated microbes. The microbe communities identified by raw abundance could not be overlapped with the correlation of reduced signature weights and function (Fig. 3B). We confirmed that Escherichia coli, Enterobacter cloacae complex, Klebsiella pneumoniae, Pseudomonas aeruginosa, and Staphylococcus aureus are highly associated with COVID-19, although some, especially those that are Gram-negative, have been found to be commonly hospital-acquired in many contexts (Asmarawati, Rosyid et al. 2021, Baskaran, Lawrence et al. 2021). While Rothia M., Fusobacterium periodonticum, Prevotella melaninogenica, and Haemophilus parainfluenzae, as well-known pathogens, were also found to link with COVID-19.

**Figure 3.**
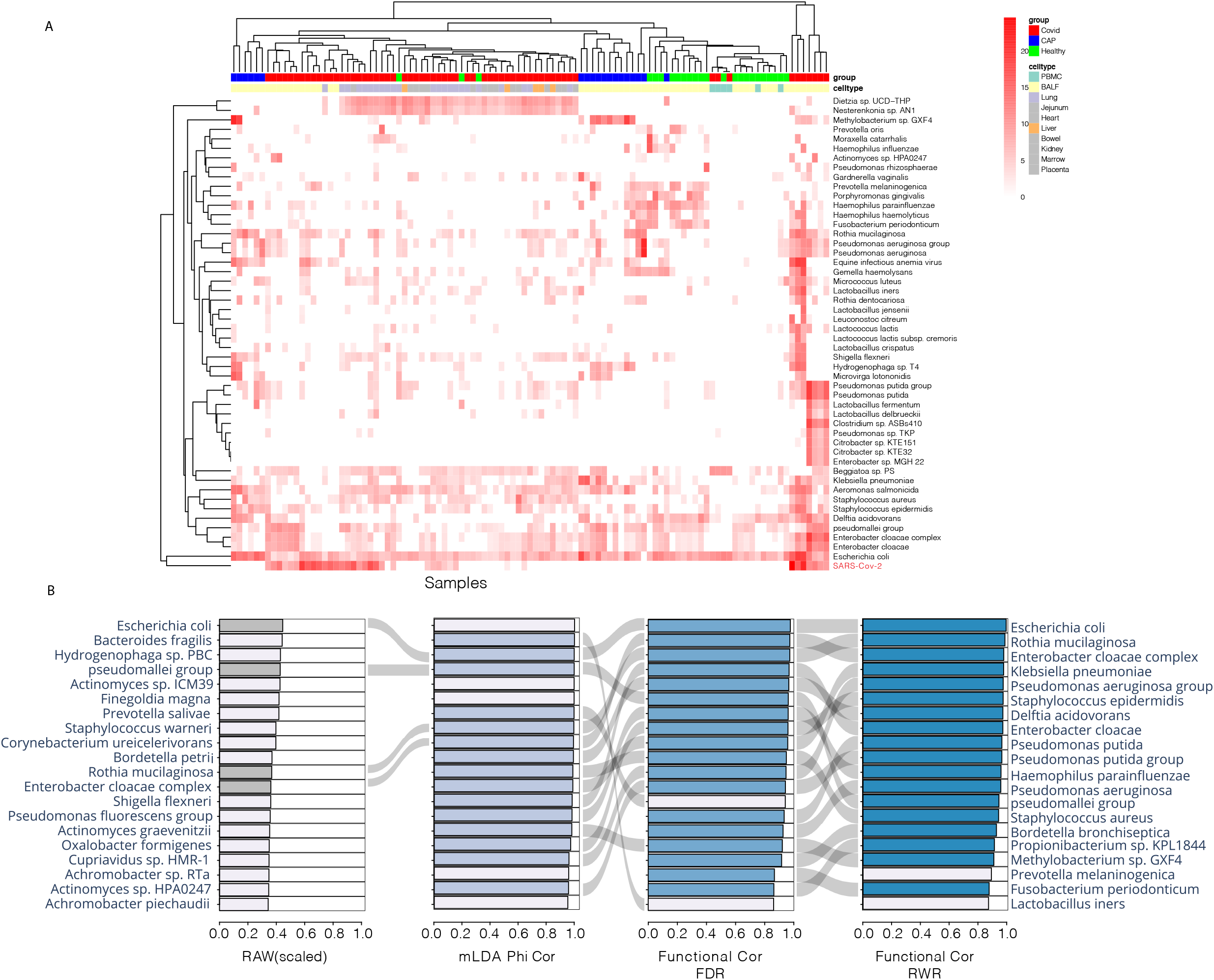
Microbe co-infection network analysis. A. Heatmap of the most abundant microbes including SARS-CoV-2. B) Co-infection of microbes with COVID-19 based on different metrics: raw abundance scaled, correlation of microbe to topic weight, correlation of interacting gene profile based on FDR, correlation of interacting gene profile based on RWR.

As shown in Figure 4, these COVID-19-associated microbes have with similar interaction profiles in the COVID-19 patient group (Fig. 4A). but they demonstrate different interacting patterns in the healthy patient group (Fig. 4B). Rothia M., Fusobacterium periodonticum, Prevotella melaninogenica, and Haemophilus parainfluenzae show significant composition changes in COVID-19 patients compared with healthy people in bronchoalveolar lavage fluid (BALF) (Fig. 4C).

**Figure 4.**
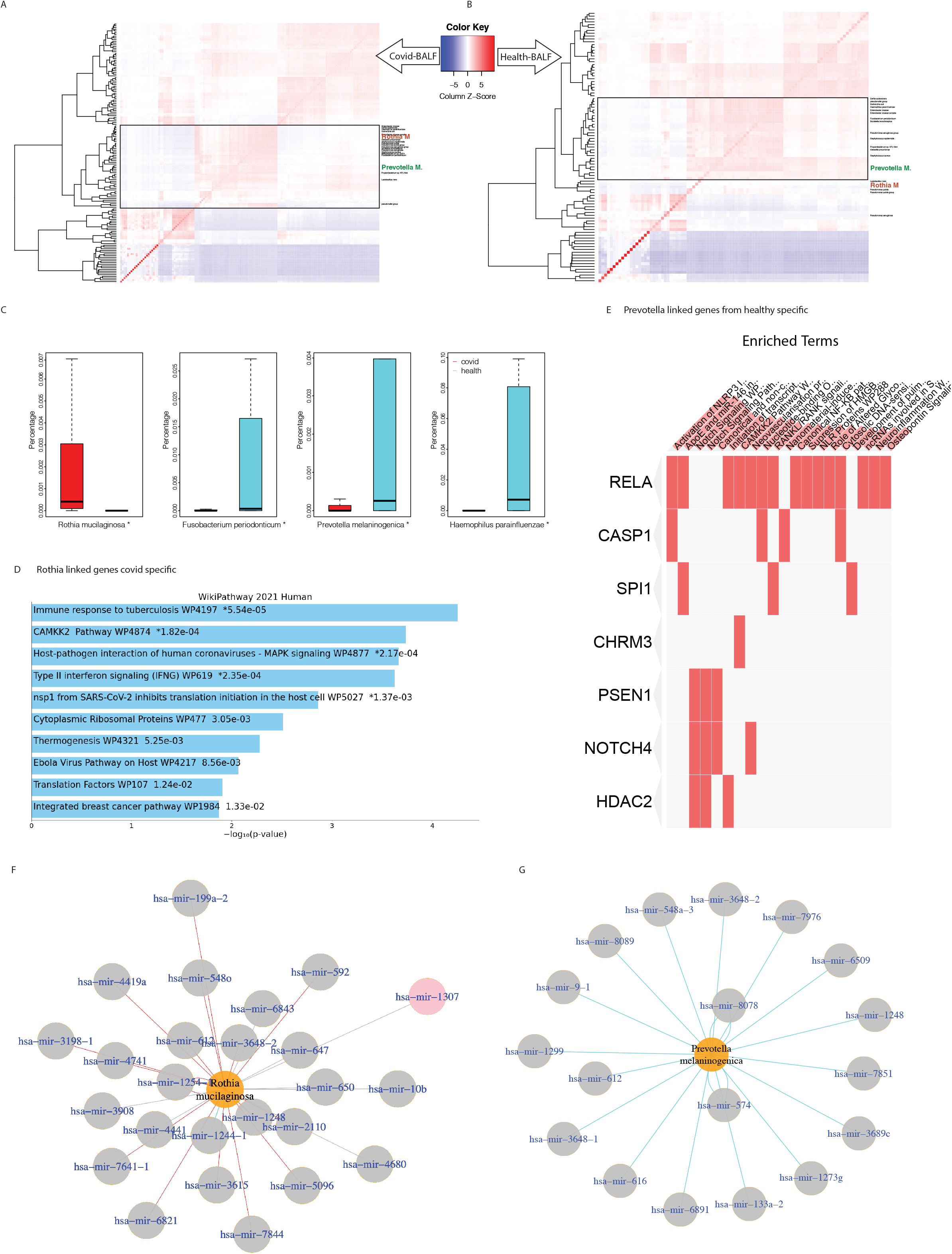
Microbe functional associations in the COVID-19 patient group. A) Heatmap of SARS-CoV-2-microbe clusters in the COVID-19 patient group. B) Heatmap of co-infection microbes in healthy patients. C) Composition of top associated microbes between COVID-19 and healthy patients. D) Functional enrichment of Rothia M. COVID-19-specific linked genes. E) Functional enrichment of Prevotella M. healthy-specific genes. F) Rothia-associated miRNA.

Rothia mucilaginosa is a gram-positive coccus that occurs as part of the normal flora of the oropharynx and upper respiratory tract (Baeza Martinez, Zamora Molina et al. 2014, Yang, Liu et al. 2020), has a significantly higher relative abundance in COVID-19 patients and showed a synergic effect with COVID-19. The specific linked genes of Rothia M. in COVID-19 patients compared with healthy individuals are highly enriched in the immune response, host-pathogen interaction, and SARS-CoV-2-related gene sets (Fig. 4D). However, when comparing the linked genes of Rothia M. between the COVID patient group and the CAP group, the significant genes are more related to cell adhesion. When comparing COVID-19 with pneumonia, the cell barriers are more important for viral infections (Fig. 4s).

Fusobacterium periodonticum, Prevotella melaninogenica, and Haemophilus parainfluenzae, show a reduced relative abundance in COVID-19 patients. In particular, the significant high composition of Prevotella M. in healthy individuals also results in specific interacting genes, which are shown in Figure 4E. RELA is a subunit of the NF-Kappa B complex, which forms the RELA-RELA complex and is involved in invasion-mediated activation of IL-8 expression and regulating the IFN response during SARS-CoV-2 infection. Another important pathway that shows significant high enrichment is the Notch signaling pathway (including genes PSEN1, NOTCH4, and HDAC2), which is significantly upregulated in the lungs in COVID-19 infections (Rosa, Ahmed et al. 2021). These genes and pathways linked to Prevotella M. are also key factors for COVID-19 infection, considering the composition of Prevotella M. is significantly decreased in COVID-19 patients. Thus, Prevotella M. may potentially have an antagonistic relationship with COVID-19.

### SARS-Cov-2 links to Genes & Pathways

We compared the linkages across 10 different SARS-CoV-2 patient samples, which included BALF, bowel, heart, jejunum, kidney, liver, lung, marrow, peripheral blood mononuclear cells, and placenta. We compared host gene-microbe and microbe-miRNA linkages. In addition to common linkages, we identified signature linkages (such as gene-microbe linkages) that are found only in one specific tissue. The genes, microbes, and miRNAs from these signature linkages are clustered in Figure 6, which shows clear tissue-specific patterns, in particular for BALF and lung tissue (Fig. 5s).

**Figure 5.**
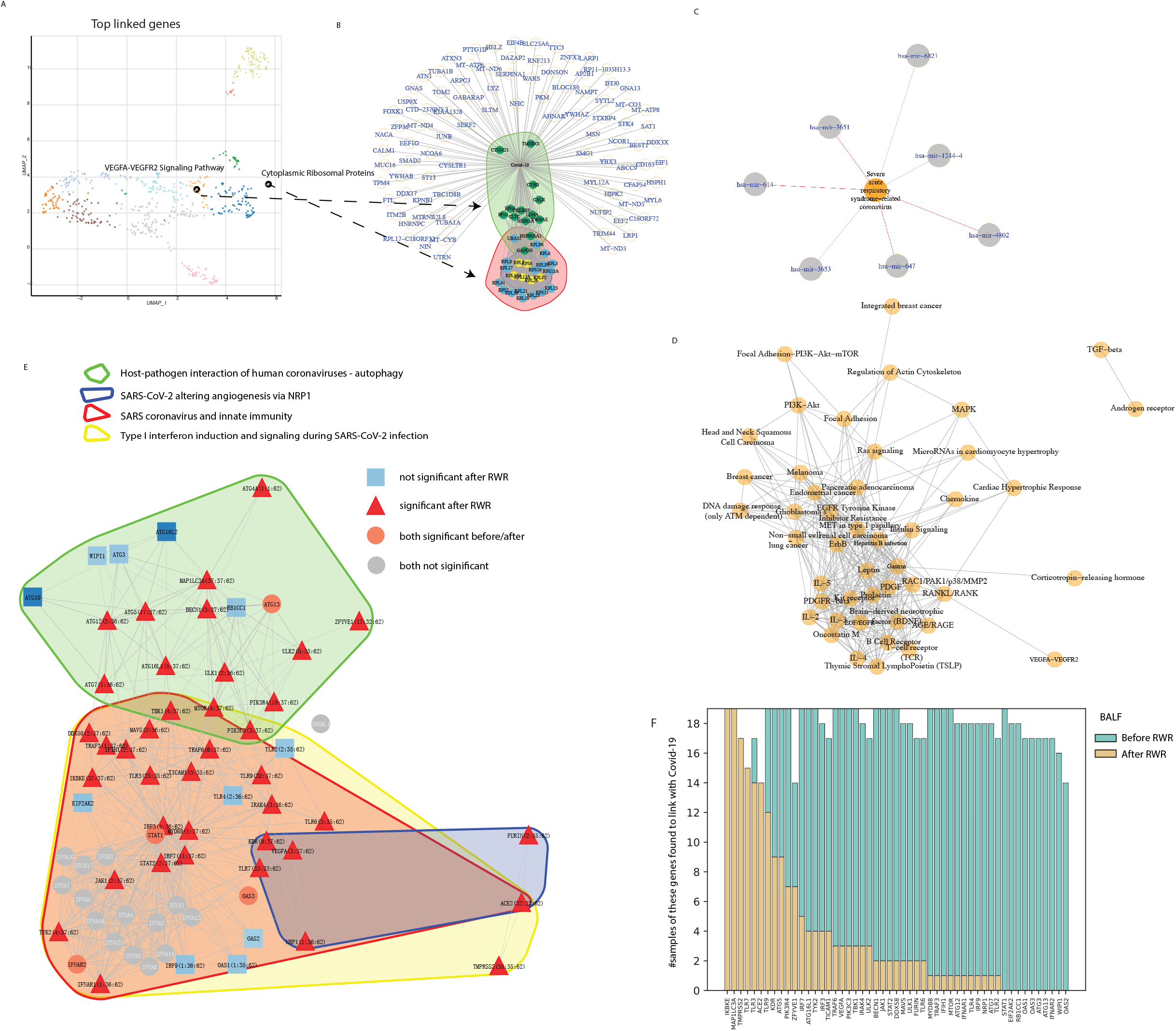
COVID-19 links. A) UMAP analysis of gene enrichment. B) Top 127 genes with highlighted enriched pathways. C) Topranked pathways from network propagation. D) COVID-19-associated miRNA. Figure 5s. Linkage comparison across different tissues. A. Gene clusters in tissue-specific gene-microbe linkages. B. Microbe clusters in tissue-specific gene-microbe linkages. C. miRNA clusters in tissue-specific miRNA-microbe linkages. D. Microbe clusters in tissue-specific miRNA-microbe linkages.

**Figure 6.**
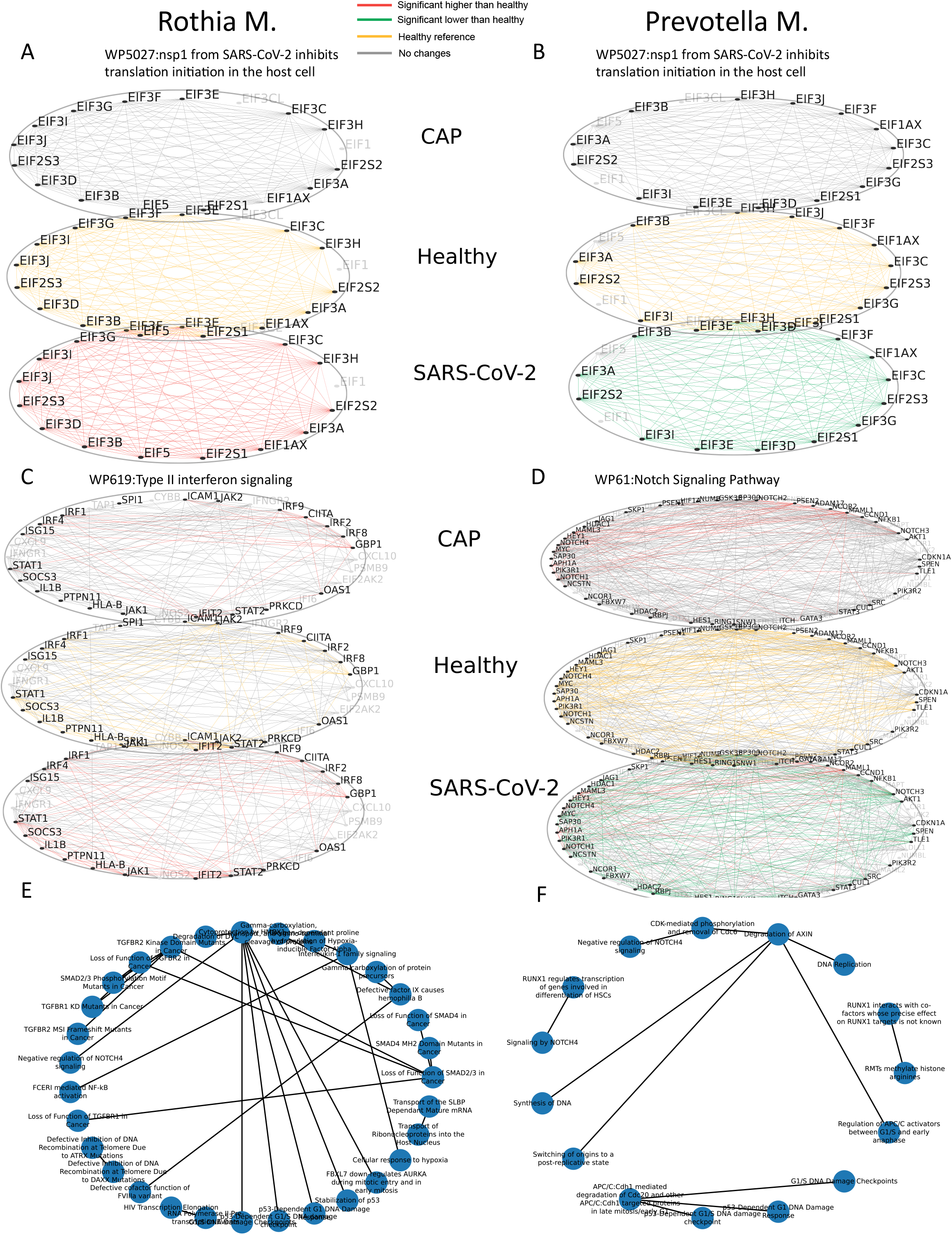
Multiple-layer comparison of co-infecting microbes potentially affect gene-gene linkages differentially. A,C) Rothia M. shows increased gene-gene linkages in the Nsp1 pathway and the type II interferon signaling pathway in COVID-19 patients; B,D) Prevotella M. shows decreased gene-gene linkages in the Nsp1 pathway and the Notch signaling pathway in COVID-19 patients; E) Rothia M. shows increased pathway-pathway connections in COVID-19 patients; F) Prevotella M. shows decreased pathway-pathway connections in COVID-19 patients

We inferred the COVID-19-specific linked genes by comparing their occurrences in the COVID-19 and healthy groups. More genes are highly represented in the COVID-19 group than those in the healthy group. The top-ranked genes and microbes associated with COVID-19 are shown in Figure 5. These genes come from many pathways. Among these genes, cytoplasmic ribosomal protein and VEGFA-VEGFR2 are significant highly associated with COVID-19 (Fig. 5 A,B). Ribosomal RNA is essential for protein synthesis in all living organisms (Schmidt, Lareau et al. 2021) and the VEGFA-VEGFR2 pathway is highly associated with viral entry.

We applied network propagation to aggregate network information. After converging the random walk with restart (RWR), we identified the top-ranked pathway with a high proportion of signature genes and gene-gene connections. The top-ranked pathway network is shown in Figure 5D. Interestingly, we consistently identified the VEGFA-VEGFR2 pathway and immune response pathway. We also identified some cancer-related pathways, which may be because COVID-19 can trigger signaling pathways that respond to cancer.

ACE2 plays important roles in the entry of COVID-19 virus to cells. Our MLCrosstalk method identified genes in the pathway related to viral entry associated with COVID-19, including IFNAR1, IFNAR2, and STAT. Our results also verified that random walk generates stable results that can recall the most biologically relevant linked targets (Fig. 5F, like ACE2, TMPRSS2) after optimization using RWR. A relatively stable number of patients with the same background showed the same gene links with COVID-19.

### Microbes & Genes together: How co-infecting microbes can differentially affect gene-gene linkages

As no SARS-CoV-2 virus can be detected in healthy patients, it is difficult to compare the effects of viral infections using SARS-CoV-2 gene linkage information. We have identified two distinct patterns of the co-infection microbes Rothia M. and Prevotella M., which have putative synergic and antagonistic effects, respectively. We studied the enrichment of the linked genes and gene-gene connections for a pathway in different individual groups (SARS-CoV-2, healthy, and CAP) from a pathway. We found that the Rothia M. linked genes have significant higher representations in the SARS-CoV-2 patient groups (edges in red) in “Nsp1 inhibits translation initiation in the host cell pathway.” By contrast, Prevotella M. showed an opposite pattern in which the SARS-CoV-2 patient group had relatively low representation of the gene-gene edges (in green). We found similar distinct patterns between Rothia and Prevotella for other pathways, especially for the immune response and signaling pathways (such as the type II interferon signaling pathway and Notch signaling pathway).

We extracted the microbe significantly linked genes for the microbes in pathways and linked two pathways together if they shared significant-gene connections according to the known host networks. We then compared the centrality of a gene-gene connection that joined one pathway with another. Rothia M. linked genes had more interconnections among SMAD2/3, TGFBR1, P53, and HMOX1-related pathways, whereas the Prevotella M. linked genes were highly enriched in the edge connections among the RUNX1, Notch, and AXIN pathways. These findings suggest that Rothia M. and Prevotella M. co-infect with SARS-Cov-2 in two very different ways.

## Discussion

Here, we present MLCrosstalk, which we specifically developed to tackle three major challenges in integrative data mining: heterogeneity and noisiness of data, multiple-type data integration, and personalized linkage identification. Using the SARS-CoV-2 dataset as an example, we demonstrate the capability of our model to capture latent patterns of multiple types of data. Moreover, we show that the sample-specific linkages inferred by MLCrosstalk have strong support of biological evidence.

MLCrosstalk extends latent Dirichlet allocation (LDA) and is capable of handling noisy and missing data. By enforcing a unified topic distribution, MLCrosstalk controls the sparsity of topics and topic components using the hyperparameters and then builds a latent representation of multiple data types within the same topic domain. Although there are alternative ways to train each type of data with the LDA model separately, a new challenge is finding topic associations between different data types for further linkage inference. In addition, multiple-type data integration can help to identify the biological underpinnings and the comprehensive linkages between different data types.

Many alternative methods can infer the overall association using large cohort datasets. For example, correlation analysis can discover the trends and associations of two features by considering the values of a set of samples. However, these methods cannot give sample-specific associations, which are important, especially for patient-specific or tissue-specific samples. MLCrosstalk infers sample-based linkages by adding the effect of sample topic distribution to adjust the association inference to consider the different host responses from personalized background/conditions.

COVID-19 is one of the most severe public health emergencies in recent years. It is thus imperative to explore the underlying biology and provide insights for the development of treatment strategies. Our MLCrosstalk method can integrate multiple data types and learn the intrinsic latent patterns in an unsupervised manner. From the most informative Topic 9, the biologically relevant genes and pathogens ranked high in the component of Topic 9. MLCrosstalk then inferred the linkages between genes and microbes and integrated pathway information using network propagations. From the results, we identified SARS-CoV-2 co-infection clusters, and Rothia M., and Prevotella M. as the representatives of two groups of microbes, with synergic and antagonistic effects, respectively. With the indepth discovery of COVID-19-related genes, we pinpointed genes in the most enriched pathways, like the VEGFA-VEGFR2 pathway, and constructed functional pathway associations based on the target genes.

## Methods

### Data collection and processing

We collected data from CAP, COVID-19, and healthy individuals. The transcriptome data were analyzed using the exceRpt pipeline. Briefly, RNA-seq reads were subjected to quality assessment using FastQC software v.0.10.1 (https://www.bioinformatics.babraham.ac.uk/projects/fastqc/) both prior to and following 3’ adapter clipping. Adapters were removed using FastX v.0.0.13 (http://hannonlab.cshl.edu/fastx_toolkit/). Identical reads were counted and collapsed to a single entry, and reads containing N’s were removed. Clipped, collapsed reads were filtered through the Univec database of common laboratory contaminants and a human ribosomal database before mapping to the human reference genome (hg19) and pre-miRNA sequences using STAR [50]. Reads that did not align were mapped against a ribosomal reference library of bacteria, fungi, and Archaea, compiled by the Ribosome Database Project [51], and then to genomes of bacteria, fungi, plants, and viruses, retrieved from GenBank [51]. In cases where RNA-seq reads aligned equally well to more than one microbe, a “last common ancestor” approach was used, and the read was assigned to the next node up the phylogenetic tree, as performed by similar algorithms [14, 52].

The gene expression, pre-miRNA and exogenous genomic, and rRNA frequency were inferred. The exogenous contents were filtered to remove the potential contamination and to keep only pathogenic microbes. The gene expression of COVID-19, CAP, and healthy individuals were quantile normalized and converted to integers with microbe and miRNA frequency.

### MLCrosstalk model

We applied and extended a topic modeling algorithm, which can integrate multiple data types. To make the continuous data work on the topic model, all of the continuous values were converted into integers and scaled to reduce computational intensity.

For any patient group or sample, *M* (*or m*) denotes the number of individuals or samples; *K* (*or k*) is the number of topics; *θ* represents the document to topic distribution, or topics; *φ* denotes the word to topic distribution, or topic component; *α*, *β* are the hyperparameters of the document to topic distribution. The input matrices include gene(*g*), microbe (*b*), and (pre)-miRNA (*r*) abundances, for which each row represents a corresponding sample and each column is a gene, microbe, or miRNA, respectively.

The generative model is shown as below:

Given the topic distribution *θ_m_* for individual *m*, the topics assigned *z_m,n_* are drawn from *θ_m_* for each gene word occurrence from N1 total genes. Then *w_m,n_* are samples from xxx

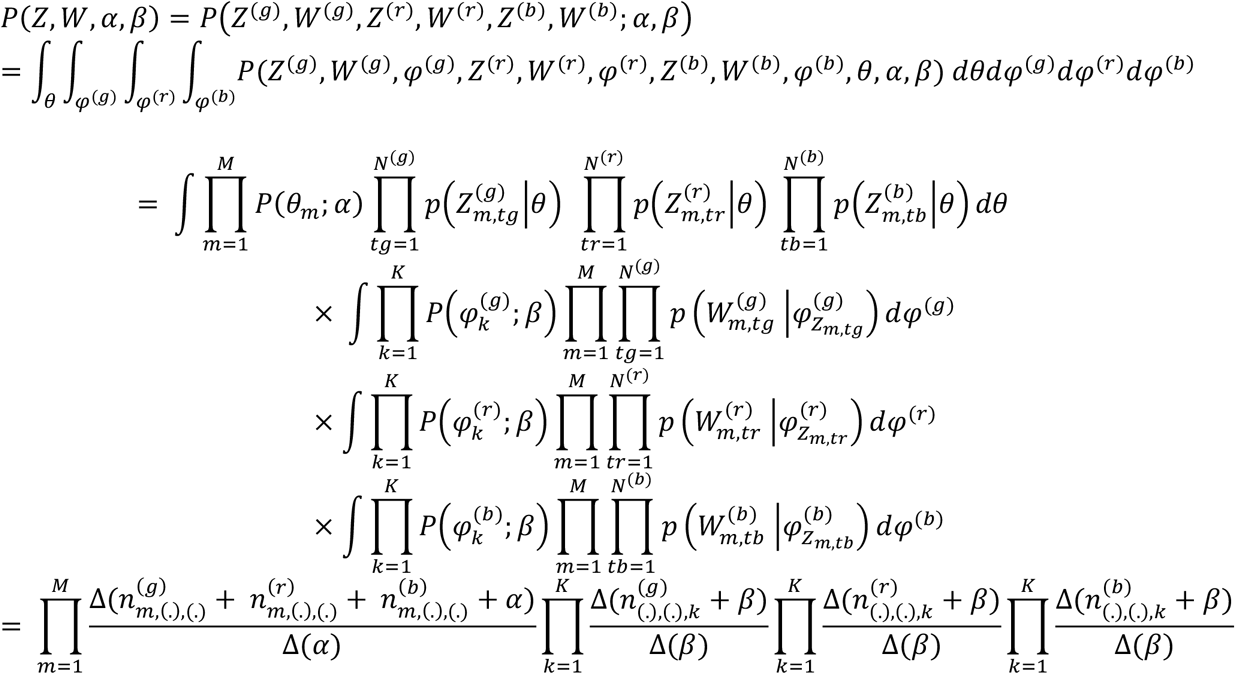

*I^G^*, *I^B^*, *I^R^* is the matrix indicator for expression and abundance, where 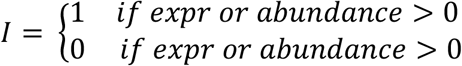 and *I* is the matrix of #word(gene, microbe or miRNA) by #sample (m).

The linkage *L_i,j_* can be defined as 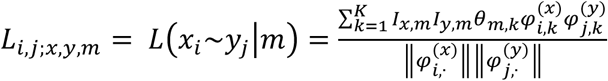, where *x*, *y* represent gene(G/g), microbe(B/b), and miR (R/r). For example, 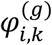 is the topic component of gene *i*, 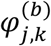 is the topic component of microbe *j*, and the linkage *L_i,j;m_*

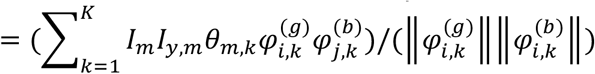

To infer a background of *L_i,j;x,y,m_*, we shuffle the *φ*^(*g*)^, *φ*^(*b*)^ for each topic *k* and then calculate the 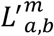 for 1,000 times and use the mean and variance to infer the one-tailed p-value. We then use the FDR adjustment to get a q-value for the inference of linkages for each sample.

### Pathway integration and curation

We used the Pathwaycommon v12 all-database version as a base, and then integrated the latest online version of KEGG (July 16, 2021) and Reactome (July 3, 2021) to output all the gene pair lists. We also combined the pathway information from WikiPathways (May 10, 2021) and gene symbols from the HUGO Gene Nomenclature Committee with the gene pair list. Finally, we obtained the gene pair list with pathway information.

### Network propagation

We generated a gene-gene interaction map based on the latest version of several PPI databases (KEGG, Reactome, and WikiPathways), in which each node represents a gene or a protein and each edge represents a gene-gene connection or PPI. Then, we applied the RWR algorithm on the network using the q-value of the microbe-gene linkage q-value as the node value.

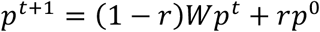

After RWR converged, we identified the top-ranked significant linked genes based on corrected q-value and pathogen link pathway score as 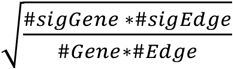

### Multiple layer analysis of changes of the host response network

Taking the results from RWR of each sample, we can analyze the linkage shown in the subnetwork for each sample. By summarizing each significant gene linkage, we can plot a subnetwork heatmap as a layer, where each layer represents a different patient group, and within a layer is the combination of all genes appearing in the RWR results, using different colors on each edge to show the frequency in which it appeared in different patient groups.

We set a cutoff value to filter significant genes in each sample, and then used them to generate a subnetwork of selected pathways for each group. By calculating how many times each edge showed up in the subnetwork after filtering, we obtained the frequency in which it appeared in each group (such as COVID-19, healthy, or CAP). Then, we set a factor to compare the edge frequency between groups. If an edge in the CAP or COVID-19 group was a factor times as large than it was in the healthy group, we marked the edge in the CAP or COVID-19 group in red, and the edge in the healthy group in yellow. If an edge in the CAP or COVID-19 group was a factor times as small than it was in the healthy group, we marked the edge in the CAP or COVID-19 group in green, and the edge in the healthy group in yellow. If a node, which represents a gene, had a related edge that was significantly larger or smaller, we marked this node in black.

## Notes

### Competing Interest Statement

The authors have declared no competing interest.

