## Supplementary figures and images for "Constructing a multiple-layer interactome for SARS-CoV-2 in the context of lung disease: Linking the virus with human genes and co-infecting microbes"

### Fig5s

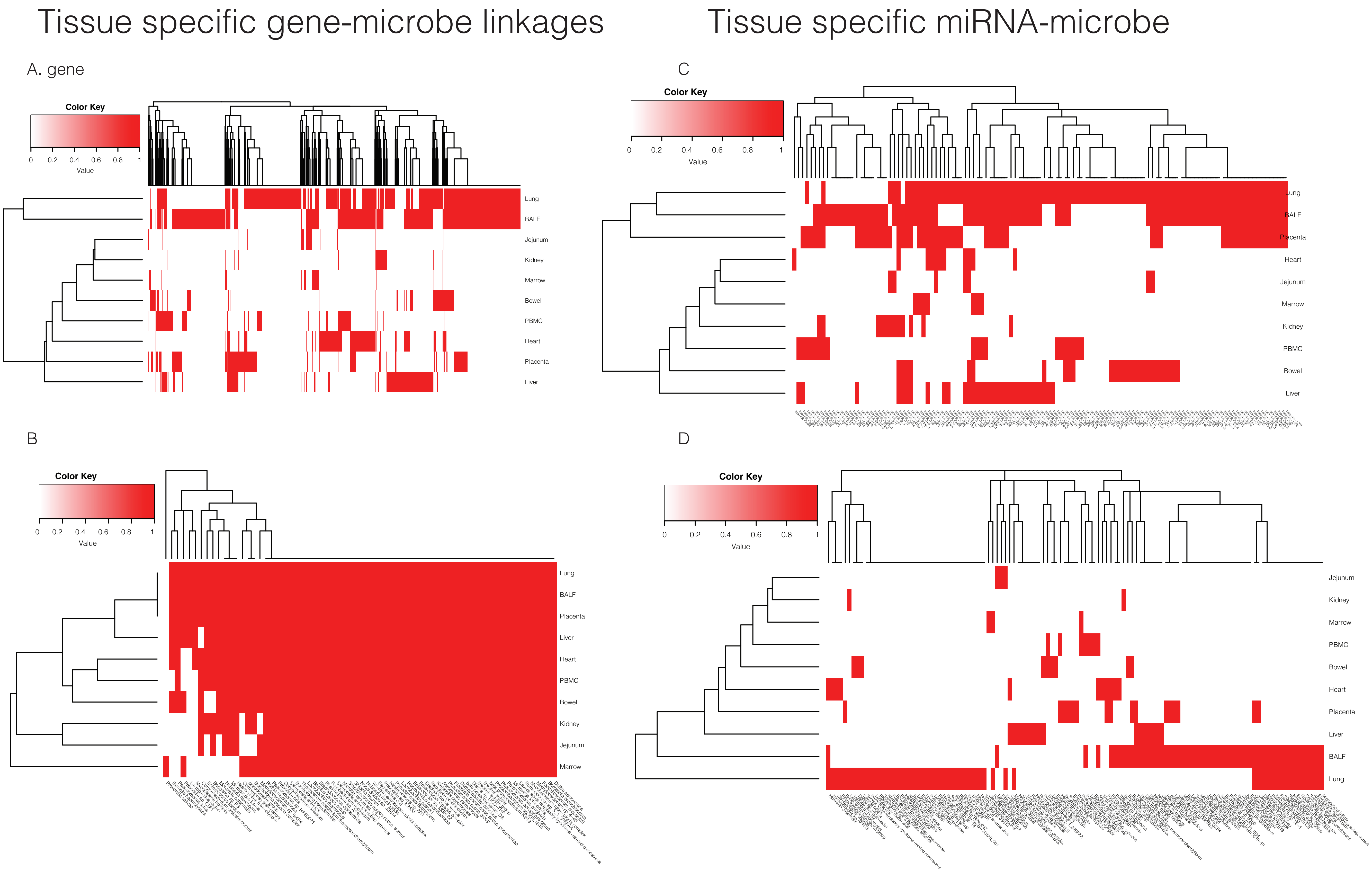
